# Relative Affinities of Protein-Cholesterol Interactions from Equilibrium Molecular Dynamics Simulations

**DOI:** 10.1101/2021.06.02.446704

**Authors:** T. Bertie Ansell, Luke Curran, Michael R. Horrell, Tanadet Pipatpolkai, Suzanne C. Letham, Wanling Song, Christian Siebold, Phillip J. Stansfeld, Mark. S. P. Sansom, Robin. A. Corey

**Affiliations:** Department of Biochemistry, University of Oxford, South Parks Road, Oxford, OX1 3QU, UK; Department of Physiology, Anatomy & Genetics, University of Oxford, South Parks Road, Oxford, OX1 3PT, UK; Sir William Dunn School of Pathology, University of Oxford, South Parks Road, Oxford, OX1 3RE, UK; Division of Structural Biology, Wellcome Centre for Human Genetics, University of Oxford, Roosevelt Drive, Oxford, OX3 7BN, UK; School of Life Sciences and Department of Chemistry, University of Warwick, Coventry, CV4 7AL, UK

## Abstract

Specific interactions of lipids with membrane proteins contribute to protein stability and function. Multiple lipid interactions surrounding a membrane protein are often identified in molecular dynamics (MD) simulations and are, increasingly, resolved in cryo-EM densities. Determining the relative importance of specific interaction sites is aided by determination of lipid binding affinities by experimental or simulation methods. Here, we develop a method for determining protein-lipid binding affinities from equilibrium coarse-grained MD simulations using binding saturation curves, designed to mimic experimental protocols. We apply this method to directly obtain affinities for cholesterol binding to multiple sites on a range of membrane proteins and compare our results with free energies obtained from density-based equilibrium methods and with potential of mean force calculations, getting good agreement with respect to the ranking of affinities for different sites. Thus, our binding saturation method provides a robust, high-throughput alternative for determining the relative consequence of individual sites seen in e.g. cryo-EM derived membrane protein structures surrounded by a plethora of ancillary lipid densities.

## Introduction

Eukaryotic integral membrane proteins participate in a range of essential cellular functions including signalling, adhesion, solute transport and ion homeostasis. Membrane proteins are inserted in a lipid bilayer, the composition of which varies between cellular compartments, metabolic state and intramembrane localisation^1,2^. Specific interactions of lipids with proteins have been observed both experimentally and in molecular dynamics (MD) simulations^3–5^ and can alter protein functionality by e.g. allosteric modulation^6–8^ or bridging protein-protein oligomerisation^9,10^.

Structural elucidation of specific protein-lipid interactions has been aided by advances in cryo-EM^11,12^. However, distinguishing the molecular identity of lipid-like densities can be challenging, and is limited to higher resolution examples^13,14^. Differentiating between phospholipid and sterol densities, is somewhat easier due to their distinct shapes. In mammalian cell membranes the most abundant sterol is cholesterol, whereas in yeast and plant cell membranes it is ergosterol and phytosterol respectively^15^. Cholesterol is typically present at concentrations of 30-40%^16,17^ although this may vary across different regions of the membrane, and is higher in sphingolipid enriched areas^18^. Cholesterol has been shown to bind and modulate a broad range of membrane proteins including G-protein coupled receptors (GPCRs), ion channels and solute transporters^6,19–23^. Recent cryo-EM structures have revealed a number of sterol-like densities surrounding protein transmembrane domains (TMD). In these instances, the bound density is either cholesterol, co-purified from the native bilayer^24–26^, or may correspond to cholesterol derivatives such as cholesterol-hemisuccinate (CHS), which are added during purification^27,28^. Often multiple cholesterol binding sites are observed within the same structure^25^. For example a recent structure of the serotonin receptor, 5-HT_1A_ (Protein Databank (PDB) ID: 7E2X), revealed 10 cholesterol molecules surrounding the TMD, including one partially buried cholesterol adjacent to the orthosteric ligand pocket^29^. There is therefore a clear need to understand and characterise the relative affinities of multiple cholesterol binding sites on the same protein. However, this remains experimentally challenging, and there is a paucity of *quantitative* experimental biophysical data for cholesterol binding to e.g. GPCRs^30,31^ and other membrane proteins.

Equilibrium MD simulations have been used extensively to expand on the information provided from structural analyses, study protein-lipid interaction patterns and obtain detailed insight into specific binding sites^4,32,33^. In addition, biased-sampling simulations, such as potential of mean force (PMF) calculations, free energy perturbation, and metadynamics simulations have been used to obtain lipid binding free energies, supplementing available experimental data on lipid binding affinities^34^. These biased simulations are often performed subsequent to initial equilibrium MD simulations, therefore requiring additional computing resource, and an iterative process to select suitable reaction coordinates. This limits the applicability of such approaches to high-throughput, automated pipelines; for example MemProtMD^35,36^. To circumvent these limitations, efforts have been made to derive protein-lipid binding affinities directly from equilibrium MD simulations. These have the advantage that multiple lipid sites can be simultaneously examined, such as in studies using 2D density distributions of cholesterol surrounding the A_2A_ and/or β_2_ adrenergic receptors, taken from either atomistic^37^ or coarse-grained (CG) MD simulations^38^. Additionally, complex lipid interaction profiles^32^ can be more readily determined, such as applied in a ‘density-threshold’ approach with the nicotinic acetylcholine receptor in a mixed lipid environment^39^. This can also be achieved with biased simulations, but requires additional simulations for each lipid species studied^40^. However, it remains unclear how accurate equilibrium methods are for obtaining binding affinities, and whether full convergence is feasible within the limits of current MD simulations.

Here, we present a method for obtaining apparent dissociation constants (K_d_^app^) directly from equilibrium MD simulations. We apply this method to rank the strength of binding sites for cholesterol on three representative membrane proteins: an ATP-dependent pump (P-glycoprotein; P-gp; see below for further details), a sterol receptor/transporter protein (Patched1; PTCH1), and a member of the TRP-family of ion channels (Polycystin-2; PC2) (Fig. 1). In particular, we investigate whether the site rankings derived from this approach are comparable with existing equilibrium and non-equilibrium methods. We also study whether these differences are maintained in the presence of higher (i.e. physiological) membrane concentrations of cholesterol. We illustrate the utility of our robust method for determining the relative affinities of multiple cholesterol sites on a membrane protein via its application to the serotonin receptor (5-HT_1A_), a GPCR structure recently determined by cryo-EM with 10 cholesterol molecules bound^29^.

**Figure 1:**
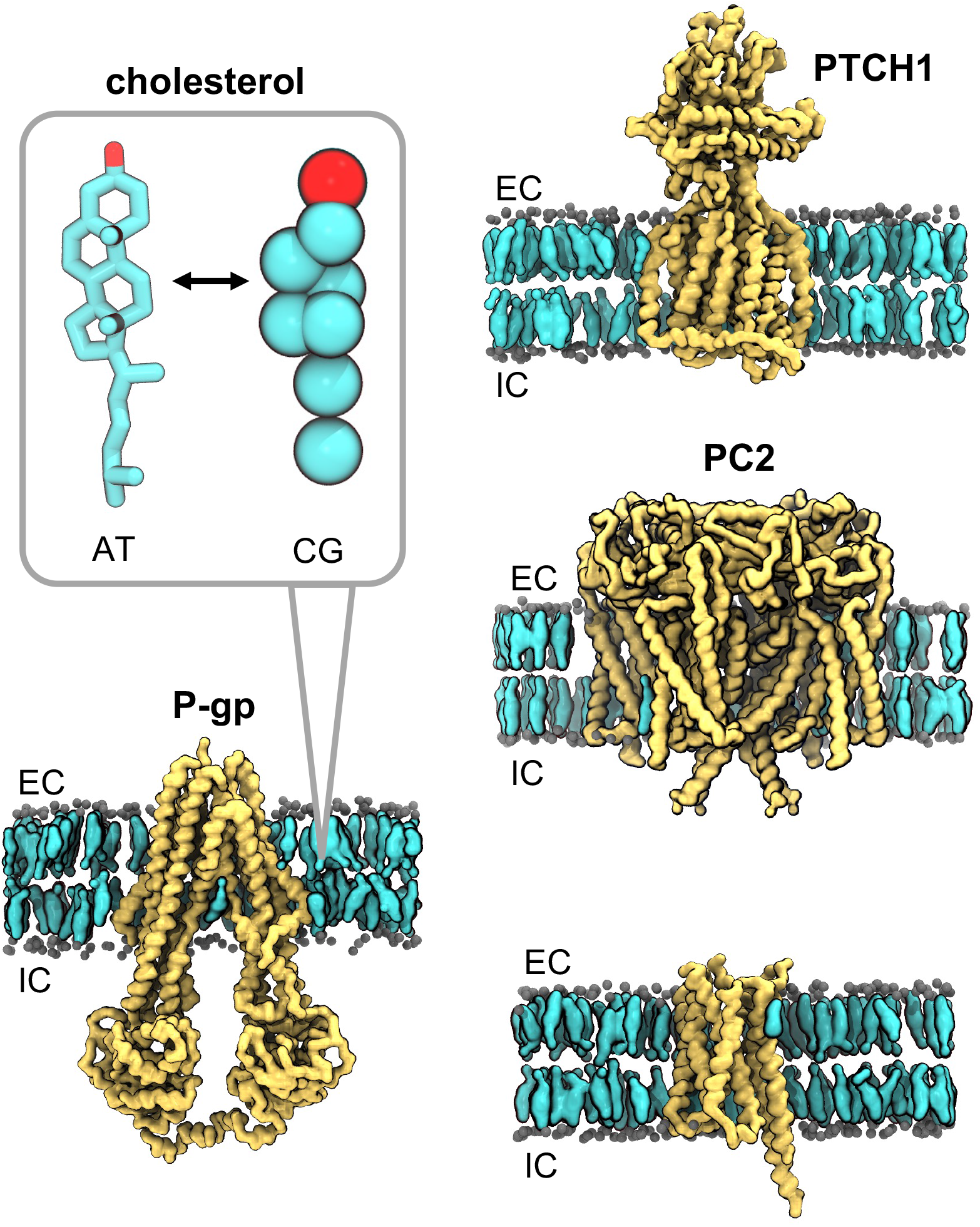
Membrane proteins which bind cholesterol. Coarse-grained (CG) representations of the structures of a transporter (P-glycoprotein; P-gp; PDB ID: 7A65, subunit A), a receptor (Patched1; PTCH1; PDB ID: 6RVD, subunit A), an ion channel (human polycystin-2; PC2; PDB ID: 6T9N, subunits A-D) and a GPCR (5-hydroxytrptamine/serotonin receptor; 5-HT_1A_; PDB ID: 7E2X, subunit R), embedded in a phosphatidylcholine (PC; 60%) and cholesterol (40%) lipid bilayer. PC phosphate beads are shown as grey spheres, cholesterol is shown in quick-surf representation in cyan, and proteins are in yellow. Extracellular (EC) and intracellular (IC) leaflets are labelled. The inset shows corresponding atomistic (AT) and CG representations of cholesterol with the β_3_-hydroxyl group (equivalent to the ROH bead at CG resolution) in red.

## Methods

### Equilibrium coarse grained MD simulations

Structures of human PC2 (PDB ID 6T9N, subunit A-D)^41^, PTCH1 (PDB ID 6RVD, subunit A)^42^, P-gp (PDB ID 7A65, subunit A)^43^ and 5-HT_1A_ (PDB ID 7E2X, subunit R) were obtained from the PDB. Non-protein components were removed and loops were modelled, using MODELLER 9.20^44^, between Q296-N305 of PC2 (for each subunit) and L608-L732 PTCH1 (using a 9 residue linker as previously described^42^). Proteins were converted to CG resolution using *martinize.py*^45^ with an ElNeDyn 2.2^46^ elastic network applied (spring force constant = 500 kJ mol^−1^ nm^−2^, cut-off = 0.9 nm). For PC2 the elastic network was applied to each subunit separately.

The MARTINI2.2^47^ forcefield was used to describe all components. Proteins were embedded in a symmetric POPC/cholesterol bilayer using *insane.py*^48^ (Fig. 1). The following cholesterol concentrations were used with the remaining bilayer composed of POPC: 1%, 2.5%, 5%, 10%, 15%, 30% and 40% cholesterol. Cholesterol was modelled using the virtual site parameters^49^. *insane.py* was also used to solvate the system with MARTINI water^45^ before neutralisation and addition of ions to ~0.15 M NaCl. Each replica was independently energy minimised using the steepest-decent method and equilibrated in 2 × 100 ns steps with restraints applied to the backbone beads.

Each protein was simulated for 5 × 5 μs in each bilayer composition (7 bilayer compositions × 5 replicates = 175 μs per system) using the GROMACS 2018 and 2019 simulation packages (www.gromacs.org). A 20 fs timestep was used and periodic boundary conditions applied. Temperature was maintained at 310 K using the V-rescale thermostat^50^ and a τ_T_ coupling constant of 1.0 ps. The Parrinello-Rahman barostat^51^ was used to maintain pressure at 1 bar with a τ_P_ value of 12 ps and compressibility of 3x 10^−4^ bar^−1^. Electrostatic interactions were cut-off at 1.1 nm using the reaction-field method and Lennard Jones interactions were cut-off at 1.1. nm using the potential-shift Verlet method. Bonds were constrained to their equilibrium values using the LINCS algorithm^52^.

### Binding site identification

Interactions of cholesterol with each protein were calculated using PyLipID (github.com/wlsong/PyLipID). Cholesterol interaction occupancy is defined as the fraction of simulation time where any bead of cholesterol is in contact with any bead of a protein residue, with a 0.55 nm /1.0 nm double cut-off used to define lipid contacts. PyLipID was also used to identify cholesterol binding sites using a community analysis approach to group residues which simultaneously interact with a bound cholesterol over the course of the trajectories. This method is described in detail elsewhere^53,54^ and has been applied to a number of recent examples to characterise lipid binding sites and kinetics^55–57^. Since the residue composition of Sites A and B varied slightly with the % cholesterol present in the bilayer, we selected six residues from each site, contacts to which were maintained across all cholesterol concentrations and used these six residues in our subsequent analysis (Supplementary Fig. 1). For 5-HT_1A_ sites were defined used 6 residues in proximity to each of the 10 modelled cholesterol densities in the structure (Supplementary Table 1).

### Binding saturation curves

To define specific interactions of cholesterol with a membrane protein we calculated the mean occupancy (Equation 1) of the six selected site residues, as reported by PyLipID, across all cholesterol concentrations. F_x_ indicates the number of frames cholesterol is bound to a given residue, F_t_ is the total number of frames and n indicates the total number of residues eg: n=6 for interactions with a site. Non-specific interactions were obtained by calculating the mean occupancy of residues which, in the 40% cholesterol system, had interactions within the 30-50% range. Further details regarding definitions of specific/non-specific interactions are included in the Supplementary Material and Supplementary Fig. 2.

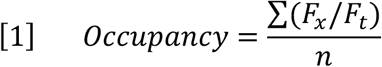

Binding saturation curves and were plotted using GraphPad Prism 9.0.2 for MacOS (www.graphpad.com). The apparent dissociation constant for cholesterol binding (K_d_^app^) was calculated by fitting the data to Equation 2, assuming site occupancies are a result of specific interactions at one site on the protein. No constraints were used in calculation of the K_d_^app^ values.

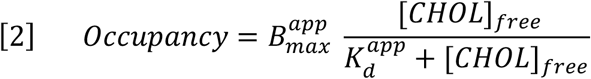

The concentration of free cholesterol ([CHOL]_free_) was derived from the mean number of cholesterol molecules > 0.8 nm from the protein surface (unbound cholesterol) as a fraction of the total number of unbound lipids (POPC and cholesterol) across simulations. Thus our computational saturation curves circumvent approximations of free and total ligand pools often used experimentally. Note that the B_max_^app^ values are not reported here (see Supplementary Material).

Convergence analyses were performed by re-running the fitting protocol with fewer simulations (Supplementary Fig. 3).

### Density analysis

We adapted a previously described method used to obtain free energy values for protein-cholesterol interactions from 2D lipid density profiles observed in simulations^37,38^. The free energy (ΔG) can be then obtained by comparing the density of cholesterol bound at a specific site (*p_site_*) to the mean lipid density in bulk (*p_bulk_*) (Equation 3). R denotes the gas constant in kJ mol^−1^ K^−1^ (8.314 × 10^−3^) and T, the temperature in kelvin.

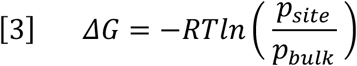

Our method utilises the same underlying approach but extends the analysis to three dimensions (density in xyz) as opposed to averaging across the bilayer normal (density in xy). Full details on processing of the density data are provided in the Supplementary Material, summarised below. Density analysis was performed using the DensityAnalysis tool implemented in MDAnalysis^58,59^ (www.mdanalysis.org) using an in-house script. Grid dimensions were fixed and the grid centre was defined as the centre of mass of the protein transmembrane domain. The bin size was 0.1 nm. Three dimensional *p_site_* and *p_bulk_* values were obtained by masking specific regions of the density array (Supplementary Fig. 4). These values were them converted directly to free energy values using Equation 3.

### Potential of mean force calculations

Setup and analysis of PMF calculations was assisted by the *pmf.py* tool (DOI: 10.5281/zenodo.3592318)^34^. CG PMF calculations were performed as described previously^34^ in bilayers containing 30% cholesterol. Briefly, a 1D reaction coordinate was generated by pulling between the cholesterol centre of mass and the backbone bead of a site residue. Windows at 0.05 nm spacing along the reaction coordinate were simulated for 1 μs each with a 1000 kJ mol^−1^ nm^−2^ umbrella pulling force used to limit cholesterol movement along this coordinate. Free energy profiles were obtained using the weighted-histogram analysis method (WHAM)^60^ implemented in GROMACS with 2000 rounds of Bayesian bootstrapping, discarding the first 200 ns of each window. Further details are provided in the Supplementary Material and convergence of free energy values is shown in Supplementary Fig. 5.

### Site membrane exposure

Membrane exposure fraction was defined as the number of lipid contacts within 0.6 nm of a bound cholesterol divided by the number of total contacts (protein and lipid) to the site cholesterol, as calculated using MDAnalysis^58,59^ across the simulations.

## Results

We set out to determine if equilibrium MD simulations are able to not only identify specific interactions of protein with lipids, but also to rank the affinities of different sites, and to evaluate how well these estimates compare to biased simulations. We also wanted to assess whether values obtained from simulations were affected by the lipid concentration in the membrane.

Using equilibrium CG MD simulations, we constructed binding saturation curves, where the total cholesterol concentration was varied, and the mean occupancy of 6 residues in each specified binding sites determined across concentrations of free cholesterol. The idea of this method was to mimic ligand binding assays used experimentally to produce binding saturation curves^61,62^.

To help with the convergence of these calculations, we chose to study the lipid cholesterol, which has been demonstrated to have relatively fast binding and/or dissociation kinetics compared to other lipids (e.g. anionic lipids such as cardiolipin and phosphatidylinositols) and hence is more amenable to sampling of multiple sites within a given simulation^33^. In addition, the thermodynamics of protein-cholesterol interactions have been extensively studied in both atomistic and CG simulations, using both biased and unbiased methods^63^. These free energy estimates therefore provide a good benchmark against which to compare our results.

Three human integral membrane proteins were selected to evaluate our analysis of cholesterol interactions: the ATP-dependant efflux pump P-glycoprotein (P-gp), the proposed sterol receptor/transporter protein Patched1 (PTCH1), and the transient receptor potential (TRP) ion channel Polycystin-2 (PC2) (Fig. 1). In each case, cholesterol has been suggested to play a role in protein function either by allosteric modulation or direct involvement in the proteins biological process.

### Comparative methods for determining cholesterol binding affinities with P-gp

Cholesterol has been shown to alter both the drug binding properties^64^ and ATP-mediated export rates^64,65^ of P-gp. In addition, P-gp localises in sphingomyelin/cholesterol enriched regions in the cell^66,67^, further supporting a role for cholesterol in modulating P-gp function.

Previously reported CG simulations of cholesterol binding to a human P-gp homology model observed cholesterol binding to multiple sites including between TM10/TM12 and TM7/TM8 which were suggested to have different free energy values (as reported by PMF calculations)^68^. The TM7/TM8 site was also observed in atomistic simulations of mouse P-gp^69^. We used PyLipID (see Methods for details) to identify two cholesterol binding sites from equilibrium simulations. Our simulations, initiated from the recently solved human P-gp structure^43^, replicated the two aforementioned cholesterol binding sites from the homology model simulations. Thus, Site A corresponds to cholesterol bound between TM10/TM12, and Site B between TM7/TM8 (Fig. 2A, Supplementary Fig. 1).

**Figure 2:**
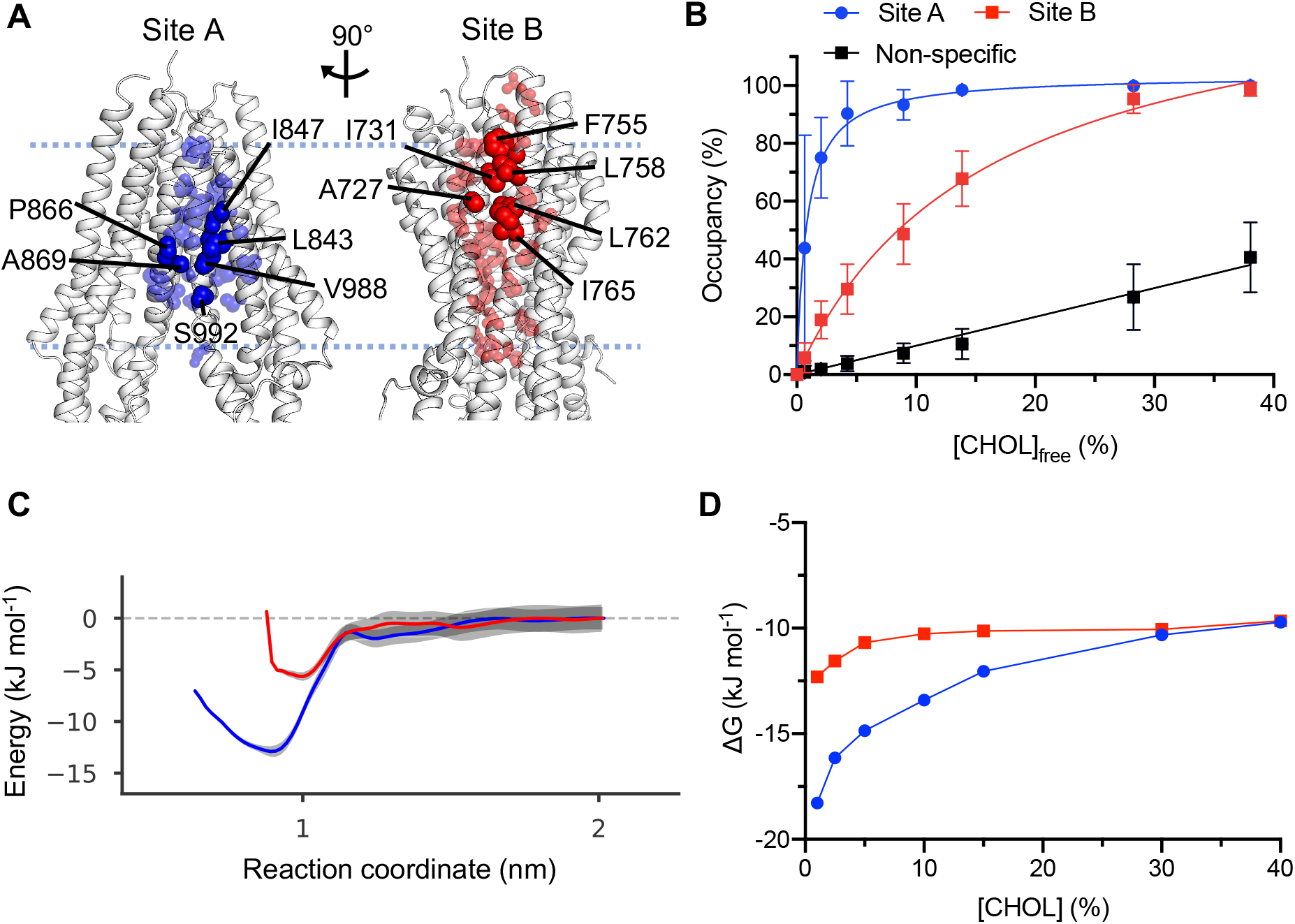
Cholesterol binding to P-gp. **A)** Cholesterol interaction Sites A (blue) and B (red), identified using equilibrium simulations (5 × 5μs at each cholesterol concentration) in PC:Chol 60:40 followed by analysis using PyLipID (github.com/wlsong/PyLipID). The sites are shown mapped onto the structure of the P-gp (7A65, subunit A) TMD. Residues involved in cholesterol interactions in the 40% cholesterol simulations are shown as spheres scaled according to cholesterol residence times. The 6 residues selected (which were conserved across all cholesterol concentrations) are labelled (opaque), whereas the remaining residues constituting the site (in 40% cholesterol) are transparent. **B)** Binding saturation curves for cholesterol binding to Site A and Site B across a range of cholesterol concentrations. Site occupancy was defined as mean occupancy of the 6 site residues in **A**. Error bars correspond to the standard deviation. Non-specific interactions were calculated from mean occupancies of specified residues with 30-50% occupancy in the 40% cholesterol simulations. **C)** Free energy landscapes from potential of mean force (PMF) calculations for Sites A and B from simulations in bilayers containing 30% cholesterol. Bootstrapping errors are shown in grey. **D)** Free energies of binding derived from probabilities of cholesterol bound at sites A (blue) or B (red) relative to the bulk probability calculated from 3D density plots of cholesterol localised surrounding P-gp.

Occupancies for both Site A and Site B increased non-linearly with cholesterol concentration, as would be anticipated for well-defined, saturable binding sites (Fig. 2B). We observe a rapid increase in Site A occupancy compared to Site B as cholesterol concentration is increased. The K_d_^app^ of P-gp Site A (K_d_^app^: 1%) is substantially lower (i.e. has a higher affinity) than for Site B (K_d_^app^: 16%). This suggests the cholesterol binding affinities of the two sites on P-gp are not equal, as is also exemplified by the variability in cholesterol affinities reported in other studies^63^.

We next performed PMF calculations to validate our observed differences in site affinities from our binding saturation method. For both sites we observe defined energetic wells at low reaction coordinate values, consistent with PMF profiles of other cholesterol binding sites^63^. We obtain a free energy well depth of −13 ± 2 kJ mol^−1^ for cholesterol binding to Site A and −6 ± 2 kJ mol^−1^ for Site B (Fig. 2C). Our PMF values are in agreement with the relative affinities of sites obtained from previous calculations on the P-gp homology model^68^. Thus, both binding saturation and PMF calculations rank the sites in the same order.

We then assessed whether density-based equilibrium free energy methods could also be used to observe quantitative differences in site binding affinities. Interestingly, for our density analysis, despite a strong difference at very low cholesterol, our free energy values for Site A and Site B converge at approximately −10 kJ mol^−1^ in 40% cholesterol. This suggests some sensitivity of the method to the lipid concentration chosen for the simulation (Fig. 2D).

### Extending analysis of cholesterol affinities to other protein examples; PTCH1 and PC2

To test the applicability of our methods to other membrane proteins we applied the same protocol described above in detail for P-gp to two other proteins: the receptor/transporter PTCH1 and the ion channel PC2. These are described in succession in the following section.

#### PTCH1

Recent structural studies of PTCH1 have identified multiple sterol binding sites on the TMD and bound within the ECD^70–74^. In addition, novel biochemical and CRISPR-based assays suggest PTCH1 alters the abundance of accessible cholesterol^73,75^, which collectively has led to the growing consensus that PTCH1 may function as a cholesterol transporter^76^.

For PTCH1, cholesterol binding sites were selected because sterol-like densities have been observed in proximity to both sites in cryo-EM structures^71,74^ (Fig. 3A, Supplementary Fig. 1). Site A is localised within a structurally conserved domain formed by TM2-6 called the sterol-binding domain (SSD). Site B is situated between TM7/TM12 of PTCH1. In addition to the observed structural densities, both sites are situated at the exit points of tunnels extending through the ECD, characterised in previous atomistic simulations, and are therefore suggested to form local cholesterol binding sites for coordination of transport between the ECD and membrane^42^.

**Figure 3:**
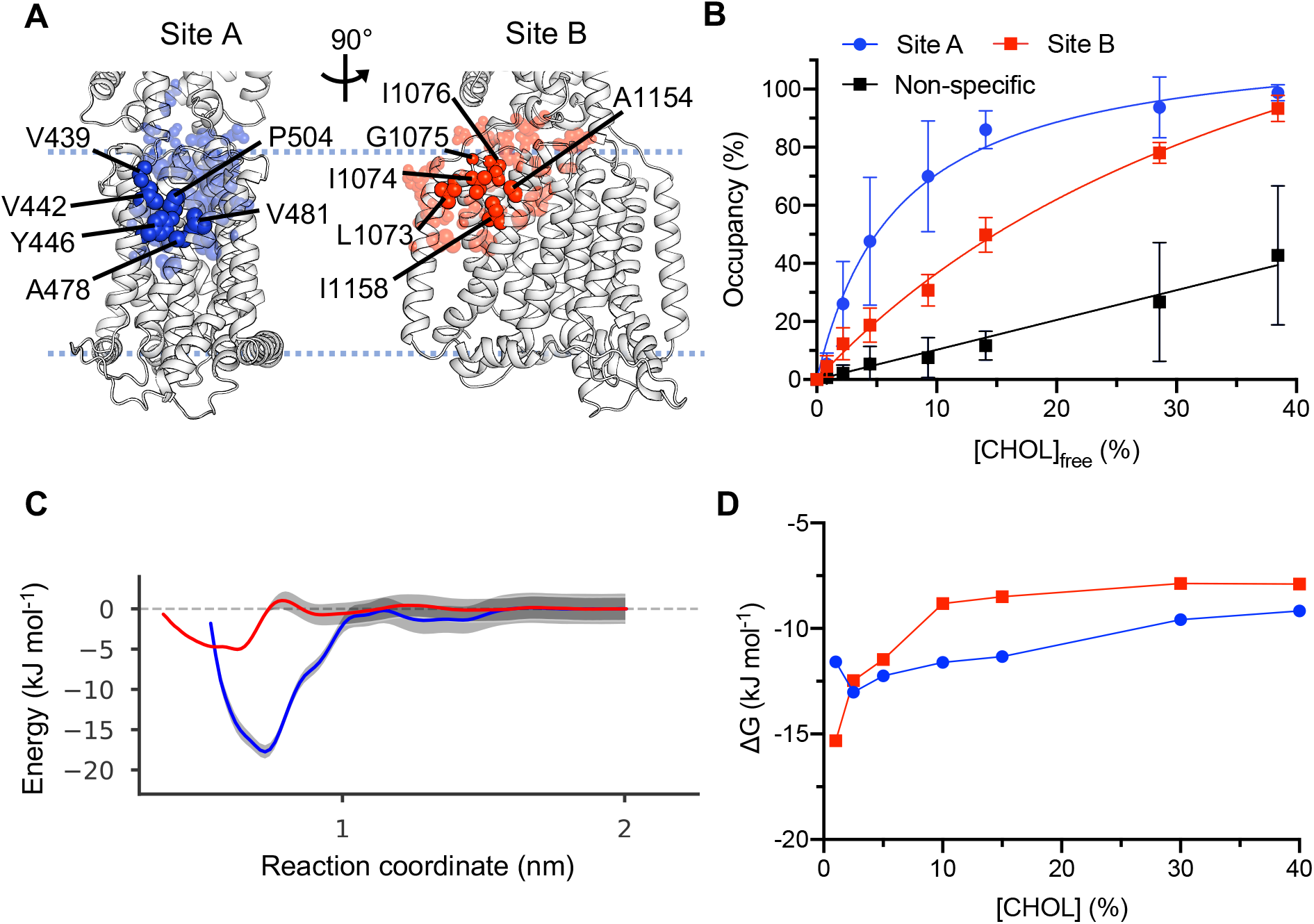
Cholesterol binding to PTCH1. As in Fig. 2 for PTCH1 (6RVD, subunit A). A) Residues comprising cholesterol interaction Site A (blue) and Site B (red) on PTCH1. B) Binding saturation curve for cholesterol binding to Site A and Site B as the concentration of cholesterol is varied. C) Free energy profiles from PMF calculations for cholesterol binding to Site A and Site B on PTCH1. D) Free energies derived from the probability of cholesterol binding to sites relative to in bulk, obtained using a density-based approach. For full methodological details see Figure 2.

We again see a strong difference in the binding saturation curves for Site A (K_d_^app^: 7%) and Site B (K_d_^app^: 46%) (Fig. 3B), suggesting that Site A has a far higher apparent affinity than Site B, although the difference between sites was somewhat less than seen in P-gp.

This difference is also reflected in our PMF calculations from which we obtain free energy well depths of −18 ± 3 kJ mol^−1^ for cholesterol binding to Site A and −6 ± 1 kJ mol^−1^ for Site B (Fig. 3C). This suggests that, not only can we obtain qualitative agreement between the ranking of site affinities using binding saturation curves and density analysis compared to PMFs, but that the magnitudes can be compared between proteins and appear to reflect genuine differences in site affinities.

These differences are reflected in our 3D density analysis, albeit with a muted difference between the sites, which at >15% cholesterol gives values of about −10 kJ mol^−1^ for Site A and −8 kJ mol^−1^ (40% cholesterol) for Site B (Fig. 3D).

#### PC2

A combined cryo-EM and MD study of PC2 identified cholesterol-like density located between the voltage-sensing-like domain (VSLD) and the pore helices, which coincided with a cholesterol binding site seen in CG simulations^41^. In addition, both PC2 and PTCH1 localise within the primary cilium of cells where levels of accessible cholesterol are regulated and where cholesterol has been shown to play roles in initiating intracellular signalling pathways^77^.

As before, we identified cholesterol binding sites on PC2 and constructed binding saturation curves. Site A on TM3/TM4 (Fig. 4A, Supplementary Fig. 1) has previously been identified from a combined structural and simulation study^41^. Site B is on the interface of TM1/TM4. We again see differences in site affinity between Site A (K_d_^app^: 11%) and Site B (K_d_^app^: 49%) from our saturation curves (Fig. 4B). The K_d_^app^ for Site A is higher than observed for P-gp or PTCH1.

**Figure 4:**
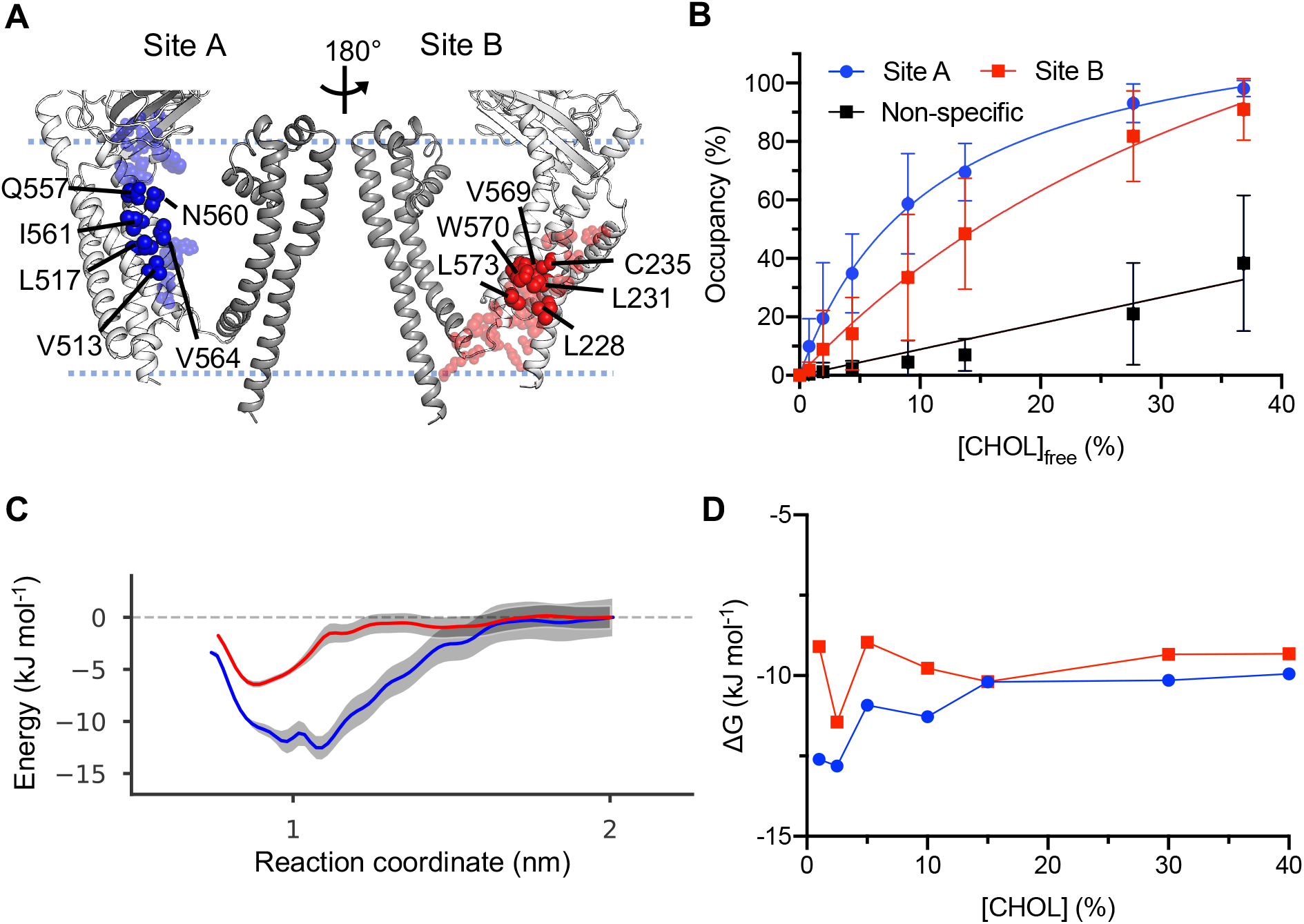
Cholesterol binding to PC2. As in Fig. 2 for PC2 (6T9N, subunits A-D). For clarify, only subunit A of the PC2 homotetramer is shown in **A**, with the pore-lining helices of PC2 in darker grey compared to the voltage-like sensing domain (VLSD). A) Residues comprising cholesterol interaction Site A (blue) and Site B (red) on PC2. B) Binding saturation curve for cholesterol binding to Site A and Site B as the concentration of cholesterol is varied. C) Free energy profiles from PMF calculations for cholesterol binding to Site A and Site B on PC2. D) Free energies derived from the probability of cholesterol binding to sites relative to in bulk, obtained using a density-based approach. For full methodological details see Figure 2.

From PMF calculations we obtain a free energy well depth of −13 ± 3 kJ mol^−1^ for cholesterol binding to Site A, in agreement with a previously reported value of −12 ± 3 kJ mol^−1^ for cholesterol binding to this site obtain by a similar method^41^ (Fig. 4C). For Site B we obtain a free energy value of −7 ± 1 kJ mol^−1^, consistent with differences in site affinities from the saturation curves.

Finally, from the density analysis we observe stabilisation of the free energy values at > 15% cholesterol, corresponding to approximately −10 kJ mol^−1^ and −9 kJ mol^−1^ for Sites A and Site B respectively (Fig. 4D).

### Affinities of multiple cholesterol sites on one protein

We sought to assess whether the binding saturation method could be applied to a membrane protein with several bound cholesterol molecules, exploiting the methods ability to obtain multiple K_d_^app^s from the same simulation dataset. For this we chose a recent structure of the 5-HT_1A_ GPCR, determined in complex with 10 cholesterol molecules^29^. Here, we used the structurally observed cholesterol densities to define the position of the binding sites (site IDs as numbered in ^29^, Supplementary Table 1) and constructed binding saturation curves for each site.

We observe saturable binding curves for all 10 sites, validating the position of modelled cholesterol densities in the 5-HT_1A_ structure^29^ (Fig. 5A). All bar one site (S_3_) had K_d_^app^s ranging between 4-9%, similar to Site A K_d_^app^s for P-gp, PTCH1 and PC2. These binding sites could be further separated into two subcategories; ‘strong’ sites with K_d_^app^s of 4-5% (S_2_, S_4_, S_7_, S_8_, S_11_) (Fig. 5B, blue, Supplementary Table 1) and ‘moderate’ sites with K_d_^app^s of 8-9% (S_1_, S_5_, S_9_, S_12_) (Fig. 5B, lime) which had distinct binding saturation profiles (Fig. 5A). The remaining site (S_3_) (Fig. 5B, orange) had an affinity (K_d_^app^ = 20%) relative to Sites A and B of P-gp, PTCH1 and PC2, suggestive of a ‘medium’ affinity site. Thus, using a single set of simulations, we are able to rank the respective affinities of the 10 cholesterol molecules bound to 5-HT_1A_.

**Figure 5:**
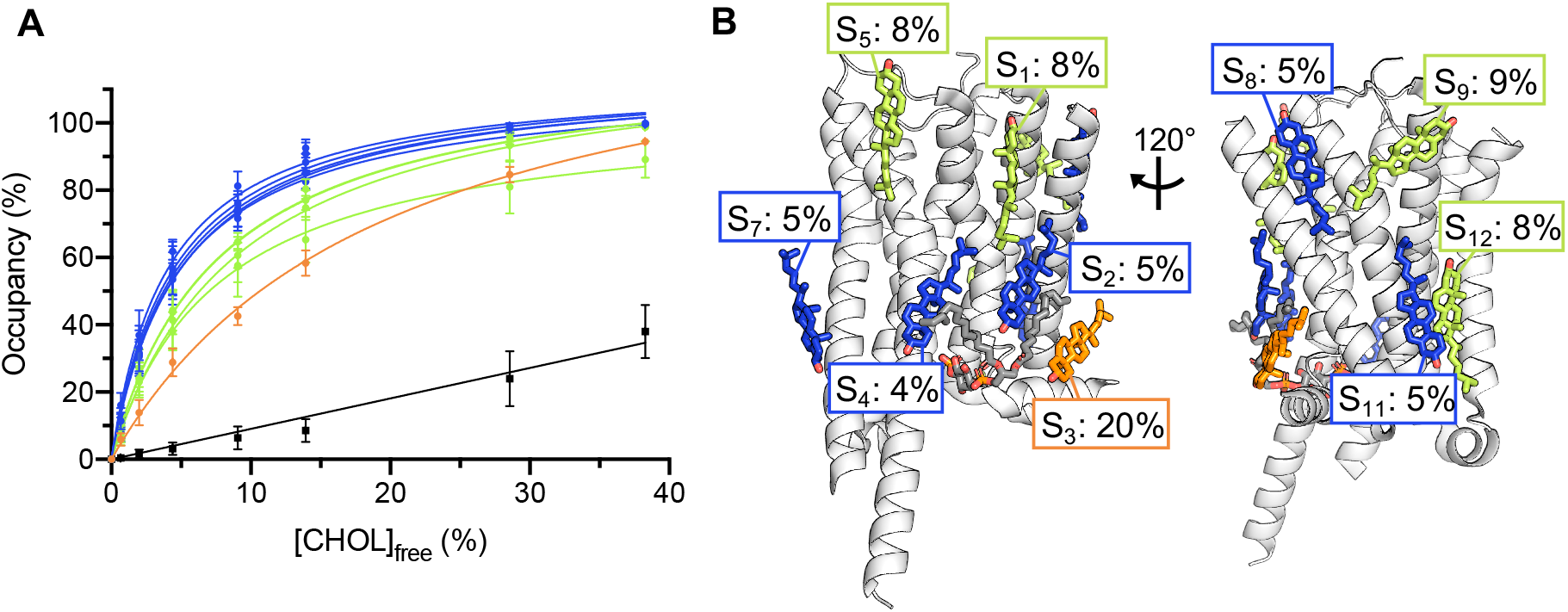
The binding saturation method as applied to 10 cholesterol sites on 5-HT_1A_. **A)** Binding saturation curves for 10 cholesterol binding sites on 5-HT_1A_ (7E2X, subunit R). Site occupancies were obtained from the mean occupancy of 6 residues in proximity to the modelled cholesterols (Supplementary Table 1), as obtained using PyLipID. Sites are coloured according to the relative strength as given by the obtained K_d_^app^ values (‘strong’, blue; ‘moderate’, lime; ‘medium’, orange). **B)** Structure of apo 5-HT_1A_ used to obtain the binding saturation curves in **A**. Cholesterol molecules are shown as sticks coloured according to the relative site affinity (see **A**) and K_d_^app^ values for each site indicated. Site IDs (S_1-5_, S_7-9_, S_11-12_) correspond to those in Fig. 2f of Xu *et al*., Nature, (2021)^29^. The modelled phosphatidylinositol is shown in grey stick representation for reference.

## Discussion

Increasingly, structures and simulations reveal a range of lipids bound to sites on the TMDs of membrane proteins^25,29,32^. Nevertheless, challenges with structural interpretation prevail when attempting to assign meaning to bound lipids in a biological context and/or for protein function. Ranking the affinity of lipid sites can aid this interpretation by establishing which sites may be more relevant/prevalent in a biological context.

We compared the affinities of two cholesterol sites on P-gp, PTCH1 and PC2 using equilibrium and biased MD simulations. Calculating the difference in apparent site free energies (ΔΔG^app^) between Sites A and B (Equation 4), reveals good agreement between the ranking of site affinities derived from PMF calculations and from our binding saturation method (Fig. 6) i.e. ΔΔG^app^ < 0 for both methods. Furthermore, the magnitude of ΔΔG^app^ between methods agrees well for P-gp and PC2, which yield ΔΔG^app^ values of −4 to −8 kJ mol^−1^. ΔΔG^app^ values for PTCH1 differ somewhat between methods which we attribute to a Site A free energy much greater than observed for cholesterol binding to other proteins by PMF calculations^63^. Thus, this suggests our binding saturation method can accurately rank the order and magnitude of site affinities when compared to robust free energy methods.

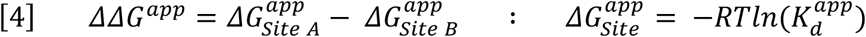

**Figure 6:**
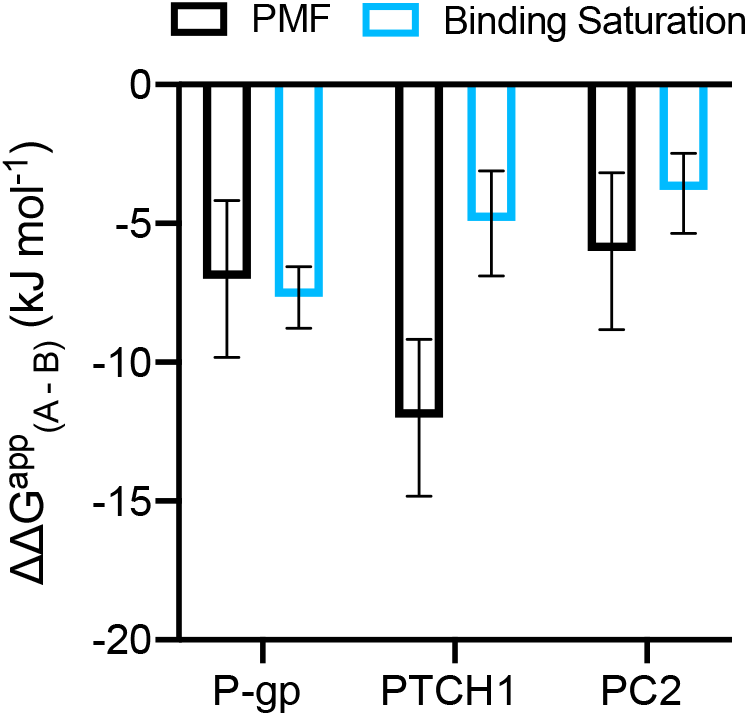
Difference in apparent free energy of binding, site A – site B. For each protein, *ΔΔG^app^* (Site A – Site B) is shown, estimated from the difference in PMF well depth (black; Fig. 2C,3C,4C) and from the difference in *ΔG^app^ = -RT ln K_d_^app^* (light blue; using K_d_^app^ values obtained from fitting the binding saturation curves in Fig. 2B/3B/4B). PMF errors were calculated in quadrature (total error = √((A_err_)^2^ + (B_err_)^2^)) since PMFs for Site A and B are independent.

Additionally, the *in silico* binding saturation method circumvents two approximations that are routinely applied in equivalent experimental procedures. Firstly, cholesterol binding occupancies are specific to the site of interest, avoiding complications created by conflating micro and macro dissociation constants and secondly, the concentration of free cholesterol can be directly calculated rather than approximating to the total cholesterol concentration. This allows for differences in site affinities to be observed over a range of physiologically relevant free cholesterol concentrations. We note that care should be taken when determining site affinities from density-based equilibrium methods as these appeared to show some sensitivity to the overall lipid concentration in the membrane (Fig. 2D/3D/4D).

The computational cost and setup ease are key considerations if we wish to investigate affinities using high throughput simulation methods applied to a wide range of membrane proteins. For the PMFs, each site on a given protein was simulated for approximately 65 × 1 μs umbrella sampling windows, in addition to the initial steered MD simulations (~0.03 μs), from which those windows were derived. Thus approximately 130 μs of CG simulation time was used in the PMF calculations to derive the two site affinities on a protein, at a single cholesterol concentration. We note that site free energy values obtained from PMFs were broadly similar at different membrane cholesterol concentrations (Supplementary Fig. 6).

For the binding saturation method, the total CG simulation time for both sites was 175 μs (5 × 5 μs at 7 lipid concentration) across all free cholesterol concentrations. The binding saturation method was surprisingly robust, with as few as 2 replicas required to reach K_d_^app^ convergence and 1 replica sufficient to observed qualitative differences in site affinities (Supplementary Fig. 3). Therefore, the total simulation time could be reduced to 35-70 μs and still yield quantitative differences in site affinities. Furthermore, the number of residues used to define the high affinity site (Site A) could be reduced from 6 to 1, reducing the amount of user input required in simulation analysis (Supplementary Fig. 7). The lower affinity site (Site B) was more sensitive to the number of site residues, as expected for weaker site binding. Setting up equilibrium simulations is more amenable to automation compared to the careful selection of reaction coordinates required in biased methods, making the former approach suitable for use in high throughput pipelines. Crucially, equilibrium methods allow site affinities to be obtained from the same simulation dataset, meaning that analyses could be extended to many sites in the same system for the same computational cost. We exemplify this here, to obtain relative affinities of 10 cholesterol sites on the 5-HT_1A_ receptor. From binding saturation curves, we can group these sites into three categories corresponding to ‘strong’, ‘moderate’ and ‘medium’ affinity sites compared to K_d_^app^s of sites on P-gp, PTCH1 and PC2 (Fig. 5). Performing ten equivalent PMF calculations would require ~650 μs of simulation time, ~4x the equilibrium CG simulation time used here. Thus, the binding saturation method is a suitable alternative for investigating site affinities, yielding tractable and accurate results with modest input required from the user.

### What dictates differences in cholesterol binding affinities?

We sought to understand whether key structural features between Site A and Site B underpin the observed differences in site affinities (Fig. 7). For P-gp and PTCH1, the membrane exposure of the bound cholesterol was lower for Site A than for Site B (Fig. 7A). This suggests that the more buried the cholesterol site is, the higher the observed affinity. That said, the degree of site exposure to the surrounding membrane was not sufficient to fully describe differences in site affinity for PC2, where both sites were similarly buried but the affinities were different. For PC2, the presence of a polar residue (Q557) in proximity to the hydroxyl (ROH) bead of cholesterol appears to enhance the affinity of cholesterol binding to Site A (Fig. 7B). Equivalent polar residues are not present in Site B (Fig. 4A). A polar residue was also present in Site A of P-gp and PTCH1 adjacent to the cholesterol ROH bead (Fig. 7B). Thus ‘strong’ cholesterol binding is enhanced by both the pocket-like nature of a binding cavity and polar residues in direct contact with the lipid headgroup.

**Figure 7:**
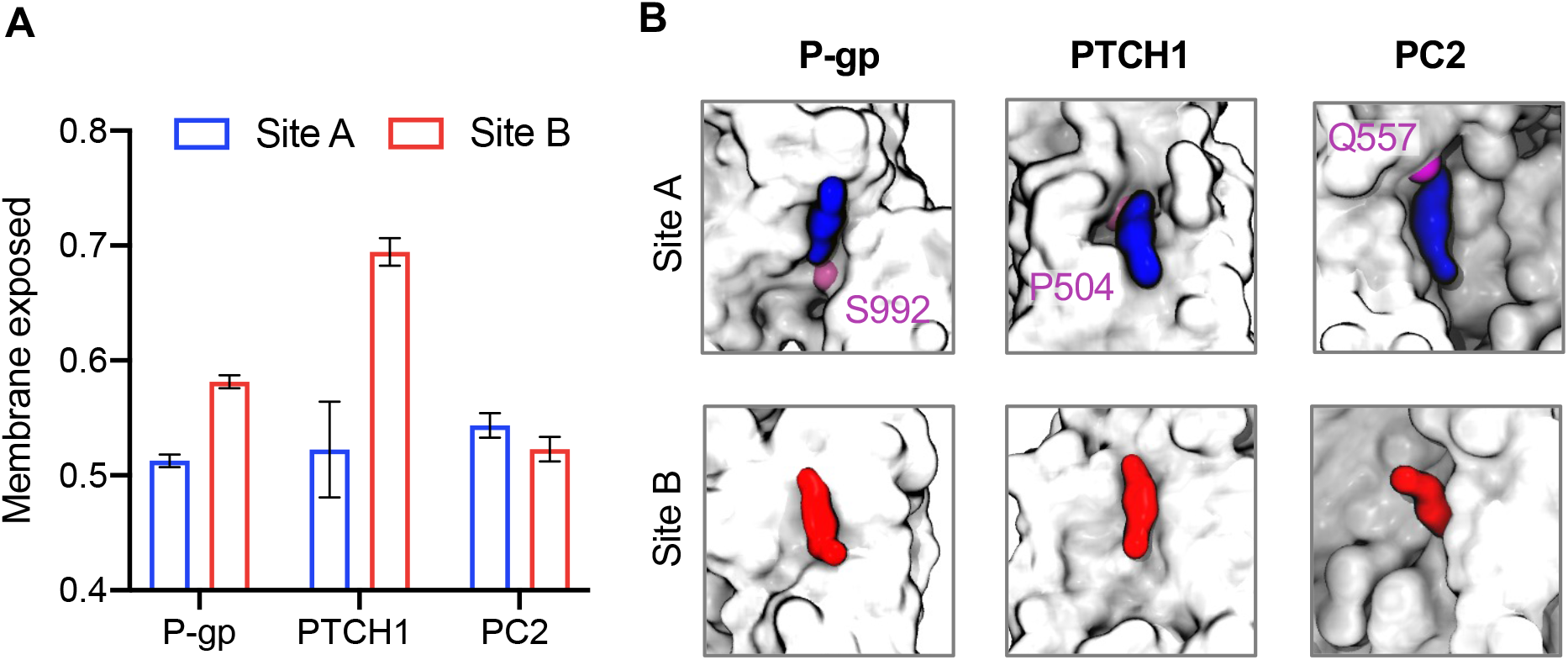
Molecular basis of observed differences in site affinities. **A)** Fractional membrane exposure of the bound Site A and Site B cholesterols for P-gp, PTCH1 and PC2 across simulations. Membrane exposure was defined as the number of lipid contacts within 0.6 nm of the bound cholesterol divided by the total number of contacts (protein and lipid) within 0.6 nm. Error bars indicate the standard error of the mean across replicates. **B)** The binding pose of cholesterol bound to Sites A (blue) and B (red) of P-gp, PTCH1 and PC2, as obtained using PyLipID from our CG simulation data. Site cholesterols are shown bound to the surface (all beads, white) of the proteins, indicating differential burial of the site cholesterols. The location of polar residues in proximity to the ROH bead of Site A cholesterols are indicated in purple.

For 5-HT_1A_, almost all sites contained polar residues in proximity to the cholesterol hydroxyl, reflected in the low K_d_^app^ values obtained from the binding saturation curves (Fig. 5, Supplementary Table 1). Higher affinity sites are more likely to persist during the relatively harsh purification and solubilisation process used to obtain membrane protein structures by cryo-EM, consistent with the high affinity of sites observed on 5-HT_1A_. One site on 5-HT_1A_ (S_1_) is proposed to stabilise the orthosteric ligand binding pocket and regulate binding of aripiprazole to the receptor^29^. We obtained a K_d_^app^ value of 8% for this site which, while reasonably high affinity, was not the strongest cholesterol binding site on 5-HT_1A_. Reduced affinity at S_1_ may assist dynamic binding/unbinding of cholesterol to this site compared to a constituently occupied, higher affinity, binding site.

Perhaps a more intriguing question is why, from a functional perspective, membrane proteins might show differences in cholesterol affinities across their surfaces. Differential site affinities on proteins could be utilised e.g. for cholesterol dependent differences in protein regulation. PC2 and PTCH1 localise to the primary cilia, where the abundance of accessible membrane cholesterol is highly regulated^78^. Changes in cilia cholesterol levels coincide with activation levels of key signalling pathways and to the subcellular localisation of PTCH1^75,79^. In addition, the abundance of membrane cholesterol within organelles increases between the endoplasmic reticulum and the plasma membrane^17^. Cholesterol binding/unbinding to sites could therefore aid in protein trafficking to its native membrane environment. For example the dynamic localisation of SNARE proteins within the trans-Golgi network and endosomes is affected by membrane cholesterol abundance, affecting SNARE recycling between membranes^80^.

One factor not considered in the study is the ability of other specific lipids to influence the affinity of a different lipid to a site. For example, the presence of PIP_2_ in a complex membrane environment enhances the affinity of PS binding to the Kir2.2 channel^53^. Additionally, we do not assess the relative affinity of different lipids binding to the same site as has been investigated in a recent study of the Kir6.2 channel^40^. Future work will seek to evaluate how changes in cholesterol concentrations influence site affinities within the context of more realistic membrane environments and assess roles lipid synergy might play in affinity modulation.

In summary, we have evaluated the binding affinities of cholesterol to two sites on a range of proteins, drawing comparisons between well-established PMF and equilibrium methods. We describe a novel binding saturation curve method for obtaining affinities from equilibrium simulations, intended to imitate experimental binding assays. This method was also applied to simultaneously probe the affinities of ten cholesterol binding sites on a protein, demonstrating how the method could be scaled for automated and/or high throughput analysis. The binding saturation method accurately ranks the order and relative magnitude of site affinities when compared to PMF calculations, and could be readily applied to study affinities of other lipid/ligand binding events.

## Acknowledgements

TBA, TP and SL are funded by Wellcome DPhil studentships. MRH is funded by a BBSRC studentship. CS is funded by Cancer Research UK (C20724/A26752) and the European Research Council (647278). RAC, WS, PJS and MSPS are funded by Wellcome (208361/Z/17/Z). Research in MSPS’s group is also supported by BBSRC (BB/R00126X/1) and EPSRC (EP/R004722/1). Research in PJS’s lab is funded by the MRC (MR/S009213/1) and BBSRC (BB/P01948X/1, BB/R002517/1BB/S003339/1). Simulations were carried out in part on ARCHER and JADE UK National Supercomputing Services, provided by HECBioSim, the UK High End Computing Consortium for Biomolecular Simulation (hecbiosim.ac.uk), which is supported by the EPSRC (EP/L000253/1). The authors thank Dr Tomas Malinauskas and Dr William J. Allen for helpful discussions when preparing the paper.

## References

1. Casares, D., Escribá, P. V. & Rosselló, C. A. Membrane lipid composition: Effect on membrane and organelle structure, function and compartmentalization and therapeutic avenues. Int. J. Mol. Sci. 20, 1–30 (2019).

2. Harayama, T. & Riezman, H. Understanding the diversity of membrane lipid composition. Nat. Rev. Mol. Cell Biol. 29, 281–296 (2018).

3. Sejdiu, B. I. & Tieleman, D. P. ProLint: a web-based framework for the automated data analysis and visualization of lipid–protein interactions. Nucleic Acids Res. gkab409, 1–7 (2021).

4. Corradi, V. et al. Lipid-Protein Interactions Are Unique Fingerprints for Membrane Proteins. Cite This ACS Cent. Sci 4, 709–717 (2018).

5. Manna, M., Nieminen, T. & Vattulainen, I. Understanding the Role of Lipids in Signaling Through Atomistic and Multiscale Simulations of Cell Membranes. Annu. Rev. Biophys. 48, 421–439 (2019).

6. Duncan, A. L., Song, W. & Sansom, M. S. P. Lipid-dependent regulation of ion channels and g protein-coupled receptors: Insights from structures and simulations. Annu. Rev. Pharmacol. Toxicol. 60, 31–50 (2020).

7. Dawaliby, R. et al. Allosteric regulation of G protein–coupled receptor activity by phospholipids. Nat. Chem. Biol. 12, 35–41 (2016).

8. Patrick, J. W. et al. Allostery revealed within lipid binding events to membrane proteins. Proc. Natl. Acad. Sci. U. S. A. 115, 2976–2981 (2018).

9. Yen, H. Y. et al. PtdIns(4,5)P2 stabilizes active states of GPCRs and enhances selectivity of G-protein coupling. Nature 559, 423–427 (2018).

10. Cong, X., Liu, Y., Liu, W., Liang, X. & Laganowsky, A. Allosteric modulation of protein-protein interactions by individual lipid binding events. Nat. Commun. 8, 1–8 (2017).

11. Yao, X., Fan, X. & Yan, N. Cryo-EM analysis of a membrane protein embedded in the liposome. Proc. Natl. Acad. Sci. U. S. A. 117, 18497–18503 (2020).

12. Cheng, Y. Membrane protein structural biology in the era of single particle cryo-EM. Curr. Opin. Struct. Biol. 52, 58–63 (2018).

13. Nakane, T. et al. Single-particle cryo-EM at atomic resolution. Nature 587, 152–156 (2020).

14. Huang, W. et al. Structure of the neurotensin receptor 1 in complex with β-arrestin 1. Nature 579, 303–308 (2020).

15. Dupont, S., Beney, L., Ferreira, T. & Gervais, P. Nature of sterols affects plasma membrane behavior and yeast survival during dehydration. Biochim. Biophys. Acta - Biomembr. 1808, 1520–1528 (2010).

16. Ray, T. K., Skipski, V. P., Barclay, M., Essner, E. & Archibald, F. M. Lipid composition of rat liver plasma membranes. J. Biol. Chem. 244, 5528–5536 (1969).

17. Van Meer, G., Voelker, D. R. & Feigenson, G. W. Membrane lipids: Where they are and how they behave. Nat. Rev. Mol. Cell Biol. 9, 112–124 (2008).

18. Sampaio, J. L. et al. Membrane lipidome of an epithelial cell line. Proc. Natl. Acad. Sci. U. S. A. 108, 1903–1907 (2011).

19. Taghon, G. J. et al. Predictable cholesterol binding sites in GPCRs lack consensus motifs. Structure 29, 1–8 (2021).

20. Lee, A. G. Interfacial Binding Sites for Cholesterol on G Protein-Coupled Receptors. Biophys. J. 116, 1586–1597 (2019).

21. Lee, A. G. Interfacial Binding Sites for Cholesterol on TRP Ion Channels. Biophys. J. 117, 2020–2033 (2019).

22. Lemel, L. et al. The ligand-bound state of a G protein-coupled receptor stabilizes the interaction of functional cholesterol molecules. J. Lipid Res. 62, 100059 (2021).

23. Manna, M. et al. Mechanism of allosteric regulation of β2 - adrenergic receptor by cholesterol. Elife 5, e18432 (2016).

24. Ruan, Z., Orozco, I. J., Du, J. & Lü, W. Structures of human pannexin 1 reveal ion pathways and mechanism of gating. Nature 584, 646–651 (2020).

25. Saotome, K. et al. Structures of the otopetrin proton channels Otop1 and Otop3. Nat. Struct. Mol. Biol. 26, 518–525 (2019).

26. Xue, J., Han, Y., Zeng, W., Wang, Y. & Jiang, Y. Structural mechanisms of gating and selectivity of human rod CNGA1 channel. Neuron 109, 1302–1313.e4 (2021).

27. Zhu, S. et al. Structure of a human synaptic GABAA receptor. Nature 559, 67–88 (2018).

28. Huang, C. S. et al. Cryo-EM structures of NPC1L1 reveal mechanisms of cholesterol transport and ezetimibe inhibition. Sci. Adv. 6, 1–12 (2020).

29. Xu, P. et al. Structural insights into the lipid and ligand regulation of serotonin receptors. Nature 592, 469–473 (2021).

30. Gater, D. L. et al. Two classes of cholesterol binding sites for the β2AR revealed by thermostability and NMR. Biophys. J. 107, 2305–2312 (2014).

31. Casiraghi, M. et al. Functional Modulation of a G Protein-Coupled Receptor Conformational Landscape in a Lipid Bilayer. J. Am. Chem. Soc. 138, 11170–11175 (2016).

32. Corradi, V. et al. Emerging Diversity In Lipid-Protein Interactions. Chem. Rev. 119, 5775–5848 (2019).

33. Grouleff, J., Irudayam, S. J., Skeby, K. K. & Schiøtt, B. The influence of cholesterol on membrane protein structure, function, and dynamics studied by molecular dynamics simulations. Biochim. Biophys. Acta - Biomembr. 1848, 1783–1795 (2015).

34. Corey, R. A., Vickery, O. N., Sansom, M. S. P. & Stansfeld, P. J. Insights into Membrane Protein–Lipid Interactions from Free Energy Calculations. J. Chem. Theory Comput. 15, 5727–5736 (2019).

35. Stansfeld, P. J. et al. MemProtMD: Automated Insertion of Membrane Protein Structures into Explicit Lipid Membranes. Structure 23, 1350–1361 (2015).

36. Newport, T. D., Sansom, M. S. P. & Stansfeld, P. J. The MemProtMD database: A resource for membrane-embedded protein structures and their lipid interactions. Nucleic Acids Res. 47, D390–D397 (2019).

37. Lee, J. Y. & Lyman, E. Predictions for cholesterol interaction sites on the A2A adenosine receptor. J. Am. Chem. Soc. 134, 16512–16515 (2012).

38. Genheden, S., Essex, J. W. & Lee, A. G. G protein coupled receptor interactions with cholesterol deep in the membrane. Biochim. Biophys. Acta - Biomembr. 1859, 268–281 (2017).

39. Sharp, L. & Brannigan, G. Spontaneous lipid binding to the nicotinic acetylcholine receptor in a native membrane. J. Chem. Phys. 154, 185102-1–185102-13 (2021).

40. Pipatpolkai, T., Corey, R. A., Proks, P., Ashcroft, F. M. & Stansfeld, P. J. Evaluating inositol phospholipid interactions with inward rectifier potassium channels and characterising their role in disease. Commun. Chem. 3, 1–10 (2020).

41. Wang, Q. et al. Lipid Interactions of a Ciliary Membrane TRP Channel: Simulation and Structural Studies of Polycystin-2. Structure 28, 169–184.e5 (2020).

42. Rudolf, A. F. et al. The morphogen Sonic hedgehog inhibits its receptor Patched by a pincer grasp mechanism. Nat. Chem. Biol. 15, 975–982 (2019).

43. Nosol, K. et al. Cryo-EM structures reveal distinct mechanisms of inhibition of the human multidrug transporter ABCB1. Proc. Natl. Acad. Sci. U. S. A. 117, 26245–26253 (2020).

44. András, F. & Andrej Ŝali. Modeller : Generation and Refinement of Homology-Based Protein Structure Models. Methods Enzymol. 374, 461–491 (2003).

45. Marrink, S. J., Risselada, H. J., Yefimov, S., Tieleman, D. P. & De Vries, A. H. The MARTINI force field: Coarse grained model for biomolecular simulations. J. Phys. Chem. B 111, 7812–7824 (2007).

46. Periole, X., Cavalli, M., Marrink, S.-J. & Ceruso, M. A. Combining an Elastic Network With a Coarse-Grained Molecular Force Field: Structure, Dynamics, and Intermolecular Recognition. J. Chem. Theory Comput. 5, 2531–2543 (2009).

47. de Jong, D. H. et al. Improved Parameters for the Martini Coarse-Grained Protein Force Field. J. Chem. Theory Comput. 9, 687–697 (2013).

48. Wassenaar, T. A., Ingólfsson, H. I., Böckmann, R. A., Tieleman, D. P. & Marrink, S. J. Computational Lipidomics with *insane* : A Versatile Tool for Generating Custom Membranes for Molecular Simulations. J. Chem. Theory Comput. 11, 2144–2155 (2015).

49. Melo, M. N., Ingólfsson, H. I. & Marrink, S. J. Parameters for Martini sterols and hopanoids based on a virtual-site description. J. Chem. Phys. 143, 243152-1–243152-12 (2015).

50. Bussi, G., Donadio, D. & Parrinello, M. Canonical sampling through velocity rescaling. J. Chem. Phys. 126, 014101 (2007).

51. Parrinello, M. & Rahman, A. Polymorphic transitions in single crystals: A new molecular dynamics method. J. Appl. Phys. 52, 7182–7190 (1981).

52. Hess Berk, Bekker, H., Berendsen, H. J. C. & Fraaije, J. G. E. M. LINCS: A linear constraint solver for molecular simulations. J. Comput. Chem. 18, 1463–1472 (1997).

53. Duncan, A. L., Corey, R. A. & Sansom, M. S. P. Defining how multiple lipid species interact with inward rectifier potassium (Kir2) channels. Proc. Natl. Acad. Sci. U. S. A. 117, 7803–7813 (2020).

54. Barbera, N., Ayee, M. A. A., Akpa, B. S. & Levitan, I. Molecular Dynamics Simulations of Kir2.2 Interactions with an Ensemble of Cholesterol Molecules. Biophys. J. 115, 1264–1280 (2018).

55. Corey, R. A. et al. Identification and assessment of cardiolipin interactions with E. coli inner membrane proteins. bioRxiv (2021). doi:doi.org/10.1101/2021.03.19.436130

56. Sefah, E. & Mertz, B. Bacterial Analogs to Cholesterol Affect Dimerization of Proteorhodopsin and Modulates Preferred Dimer Interface. J. Chem. Theory Comput. (2021). doi:10.1021/acs.jctc.0c01174

57. Ansell, T. B., Song, W. & Sansom, M. S. P. The Glycosphingolipid GM3 Modulates Conformational Dynamics of the Glucagon Receptor. Biophys. J. 119, 300–313 (2020).

58. Gowers, R. et al. MDAnalysis: A Python Package for the Rapid Analysis of Molecular Dynamics Simulations. Proc. 15th Python Sci. Conf. 98–105 (2016). doi:10.25080/majora-629e541a-00e

59. Michaud-Agrawal, N., Denning, E. J., Woolf, T. B. & Beckstein, O. MDAnalysis: A Toolkit for the Analysis of Molecular Dynamics Simulations. J. Comput. Chem. 32, 2319–2327 (2011).

60. Hub, J. S., De Groot, B. L. & Van Der Spoel, D. g_wham- A free Weighted Histogram Analysis implementation including robust error and autocorrelation estimates. J. Chem. Theory Comput. 6, 3713–3720 (2010).

61. Cabanos, C., Wang, M., Han, X. & Hansen, S. B. A Soluble Fluorescent Binding Assay Reveals PIP2 Antagonism of TREK-1 Channels. Cell Rep. 20, 1287–1294 (2017).

62. Ko, D. C., Binkley, J., Sidow, A. & Scott, M. P. The integrity of a cholesterol-binding pocket in Niemann-Pick C2 protein is necessary to control lysosome cholesterol levels. Proc. Natl. Acad. Sci. U. S. A. 100, 2518–2525 (2003).

63. Corey, R. A., Stansfeld, P. J. & Sansom, M. S. P. The energetics of protein-lipid interactions as viewed by molecular simulations. Biochem. Soc. Trans. 48, 25–37 (2020).

64. Eckford, P. D. W. & Sharom, F. J. Interaction of the P-glycoprotein multidrug efflux pump with cholesterol: Effects on ATPase activity, drug binding and transport. Biochemistry 47, 13686–13698 (2008).

65. dos Santos, S. M., Weber, C. C., Franke, C., Müller, W. E. & Eckert, G. P. Cholesterol: Coupling between membrane microenvironment and ABC transporter activity. Biochem. Biophys. Res. Commun. 354, 216–221 (2007).

66. Luker, G. D., Pica, C. M., Kumar, A. S., Covey, D. F. & Piwnica-Worms, D. Effects of cholesterol and enantiomeric cholesterol on p-glycoprotein localization and function in low-density membrane domains. Biochemistry 39, 7651–7661 (2000).

67. Orlowski, S., Martin, S. & Escargueil, A. P-glycoprotein and ‘lipid rafts’: Some ambiguous mutual relationships (floating on them, building them or meeting them by chance?). Cell. Mol. Life Sci. 63, 1038–1059 (2006).

68. Domicevica, L., Koldsø, H. & Biggin, P. C. Multiscale molecular dynamics simulations of lipid interactions with P-glycoprotein in a complex membrane. J. Mol. Graph. Model. 80, 147–156 (2018).

69. Thangapandian, S., Kapoor, K. & Tajkhorshid, E. Probing cholesterol binding and translocation in P-glycoprotein. Biochim. Biophys. Acta - Biomembr. 1862, 1–11 (2020).

70. Gong, X. et al. Structural basis for the recognition of Sonic Hedgehog by human Patched1. Science 361, eaas8935 (2018).

71. Qi, X., Schmiege, P., Coutavas, E., Wang, J. & Li, X. Structures of human Patched and its complex with native palmitoylated sonic hedgehog. Nature 560, 128–132 (2018).

72. Qi, X., Schmiege, P., Coutavas, E. & Li, X. Two Patched molecules engage distinct sites on Hedgehog yielding a signaling-competent complex. Science 362, eaas8843 (2018).

73. Zhang, Y. et al. Structural basis for cholesterol transport-like activity of the Hedgehog receptor Patched. Cell 175, 1–13 (2018).

74. Qi, C., Minin, G. Di, Vercellino, I., Wutz, A. & Korkhov, V. M. Structural basis of sterol recognition by human hedgehog receptor PTCH1. Sci. Adv. 5, 1–10 (2019).

75. Kinnebrew, M. et al. Cholesterol accessibility at the ciliary membrane controls hedgehog signaling. Elife 8e50051, 1–28 (2019).

76. Radhakrishnan, A., Rohatgi, R. & Siebold, C. Cholesterol access in cellular membranes controls Hedgehog signaling. Nat. Chem. Biol. 16, 1303–1313 (2020).

77. Luchetti, G. et al. Cholesterol activates the G-protein coupled receptor smoothened to promote hedgehog signaling. Elife 5, 1–22 (2016).

78. Nechipurenko, I. V. The Enigmatic Role of Lipids in Cilia Signaling. Front. Cell Dev. Biol. 8, 1–10 (2020).

79. Weiss, L. E., Milenkovic, L., Yoon, J., Stearns, T. & Moerner, W. E. Motional dynamics of single Patched1 molecules in cilia are controlled by Hedgehog and cholesterol. Proc. Natl. Acad. Sci. 116, 5550–5557 (2019).

80. Enrich, C., Rentero, C., Hierro, A. & Grewal, T. Role of cholesterol in SNARE-mediated trafficking on intracellular membranes. J. Cell Sci. 128, 1071–1081 (2015).

