## Supplementary information for "Relative Affinities of Protein-Cholesterol Interactions from Equilibrium Molecular Dynamics Simulations"

**Supplementary Material**

**Extended methods**

Binding saturation curves

A number of metrics were evaluated for characterising specific and non-specific cholesterol interactions with a membrane protein (Supplementary Figure 2). For specific interactions, tested metrics include i) the fraction of simulation time where lipid contacts any residue involved in site formation (total binding site occupancy), ii) mean occupancy of all residues involved in site formation (binding site mean occupancy) and iii) mean occupancy of the six selected residues conserved across all cholesterol concentrations (residue mean occupancy). Since the mean occupancy of the six selected residues was not influenced by the total number of site residues identified by PyLipID this metric is less biased than i) or ii), while still producing rational saturation curves. We therefore used the mean occupancy of the six site residues to calculate the specific occupancy at each site.

In addition, three metrics for calculating non-specific interactions were devised. These corresponded to i) mean occupancy of residues which interacted with cholesterol but were not identified as belonging to any specific site identified by PyLipID (null site), ii) mean occupancy of residues with occupancies less than the % cholesterol of the system (cholesterol threshold) and iii) mean occupancy of residues which, in the 40% cholesterol system, had occupancies in the 30-50% cholesterol range (40% cholesterol residues). All three metrics produced linear plots of occupancy vs cholesterol concentration, as expected for non-specific interactions however metrics i) and ii) were skewed by a number of residues with very low occupancies. In contrast metric iii) showed a linear increase in occupancy with a gradient of ~1 and was therefore used to define non-specific interactions.

Note that we do not report the calculated B_max_^app^ values. In an ensemble experiment, B_max_ reports on the total number of sites available, which is typically a higher value than the sites which can be bound by ligand in an experimental setup. Here, we have applied a single-molecule setup, with some sites readily occupied ca. 100% of the time within our simulation time-frames. In addition, we have a substantial background binding signal, owing to the high availability of cholesterol (many fold more abundant than a soluble ligand would be). Therefore, the B_max_^app^ values in our setup are simply reporting on the maximum occupancy for each site, plus a non-specific background binding likelihood.

Density analysis

Density analysis was performed using the DensityAnalysis tool implemented in MDAnalysis ([www.mdanalysis.org](http://www.mdanalysis.org))^1,2^ using an in-house script. DensityAnalysis was used to calculate the density of all cholesterol molecules across the simulation. The output was a 3D array of density values. The grid centre of the box was positioned at the centre of mass of the transmembrane region of the protein and grid dimensions (in x, y and z) were specified to standardise the array size of the density outputs. The dimensions of z were limited to the width of the bilayer (7 nm). The bin size was 0.1 nm. DensityAnalysis was used to create two binary masks corresponding to a) cholesterol densities > 0.8 nm away from the protein transmembrane domain backbone beads (bulk density mask) and b) cholesterol densities within 0.6 nm of any of the six identified site residues (site density mask). These masks were used to extract densities from the total cholesterol density arrays. The bulk density (*p_bulk_*) was defined as the mean density of cholesterol within the bulk density mask. The site density (*p_site_*) was defined at the mean density of the highest 100 cholesterol densities within the site density mask. For PC2 site densities were averaged across the four chains. The free energy was calculated directly from the *p_site_*/*p_bulk_* ratio using Equation 3 (main text).

Potential of mean force calculations

The pmf.py tool was used to setup and analyse PMF calculations (DOI: 10.5281/zenodo.3592318)^3^. First, a 1D reaction coordinate was defined between the centre of mass of the site cholesterol and the backbone bead of the following residues: PC2 (Site A: A563, Site B: W570), PTCH1 (Site A: P504, Site B: A1154) and P-gp (Site A: G984, Site B: F732). The cholesterol was pulled out of the binding site along this reaction coordinate using steered MD with an umbrella potential force of 1000 kJ mol^-1^ nm^-2^ and a pull rate of 0.1 nm ns^-1^. Superfluous rotation of the protein was prevented by applying 1000 kJ mol^-1^ nm^-2^ position restraints to two backbone beads: PC2 (A647 of chain A/C), PTCH1 (A1088/A1157) and P-gp (A123/A761). Frames were extracted at 0.05 nm intervals along the reaction coordinate and simulated for 1 μs each. A 1000 kJ mol^-1^ nm^-2^ umbrella pulling force was used to confine the cholesterol position in each window. The GROMACS weighted-histogram analysis method^4^ (WHAM) was used to construct free energy profiles along the reaction coordinates. The first 200 ns of each window was discarded as equilibration time and error estimated using 2000 Bayesian bootstraps.

**Supplementary Table 1:** Cholesterol binding sites on 5-HT_1A_. Site IDs correspond to numbered densities in Xu *et al*., Nature (2021), Fig. 2f ^5^.

| **Site ID** | **K_d_^app^ (%)** | **Binding Site Residues** | **Rank** |
| --- | --- | --- | --- |
| 1 | 8 | Y35, Q97, L380, A383, I384, W387 | ‘moderate’ |
| 2 | 5 | L46, C49, L394, L395, V398, F403 | ‘strong’ |
| 3 | 20 | C56, Y402, Q408, F411, K412, I415 | ‘medium’ |
| 4 | 4 | V344, K345, G348, I349, G352, L356 | ‘strong’ |
| 5 | 8 | V364, H376, M377, P378, L381, I385 | ‘moderate’ |
| 7 | 5 | L209, L212, V213, G216, F219, R220 | ‘strong’ |
| 8 | 5 | S40, G44, I47, P91, L95, V98 | ‘strong’ |
| 9 | 9 | V87, L88, M92, V107, T108, L111 | ‘moderate’ |
| 11 | 5 | V58, A62, G76, V80, L83, M84 | ‘strong’ |
| 12 | 8 | Y73, M84, P150, A154, I157, W161 | ‘moderate’ |

**References**

1.	Gowers, R. *et al.* MDAnalysis: A Python Package for the Rapid Analysis of Molecular Dynamics Simulations. *Proc. 15th Python Sci. Conf.* 98–105 (2016). doi:10.25080/majora-629e541a-00e

2.	Michaud-Agrawal, N., Denning, E. J., Woolf, T. B. & Beckstein, O. MDAnalysis: A Toolkit for the Analysis of Molecular Dynamics Simulations. *J. Comput. Chem.* **32**, 2319–2327 (2011).

3.	Corey, R. A., Vickery, O. N., Sansom, M. S. P. & Stansfeld, P. J. Insights into Membrane Protein–Lipid Interactions from Free Energy Calculations. *J. Chem. Theory Comput.* **15**, 5727–5736 (2019).

4.	Hub, J. S., De Groot, B. L. & Van Der Spoel, D. g_wham- A free Weighted Histogram Analysis implementation including robust error and autocorrelation estimates. *J. Chem. Theory Comput.* **6**, 3713–3720 (2010).

5.	Xu, P. *et al.* Structural insights into the lipid and ligand regulation of serotonin receptors. *Nature* **592**, 469–473 (2021).
