## Supplementary Figures for "Relative Affinities of Protein-Cholesterol Interactions from Equilibrium Molecular Dynamics Simulations"

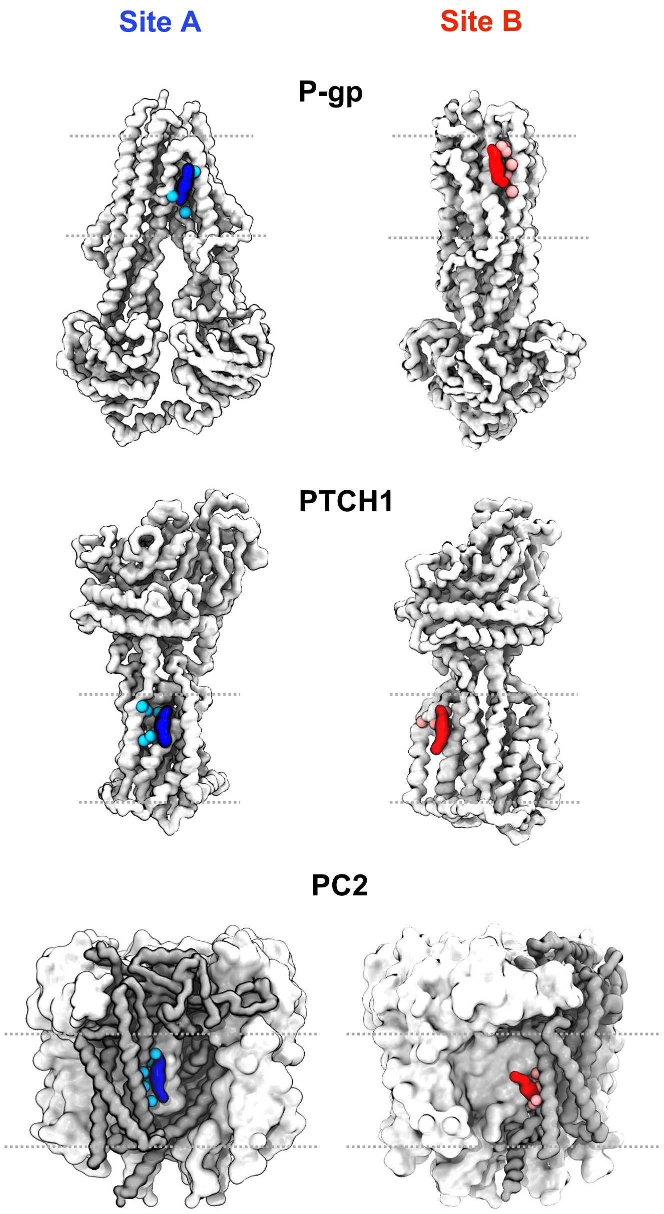


**Supplementary Figure 1: Cholesterol binding sites on membrane proteins.** Coarse-grained (CG) representations of the structures of P-gp (PDB ID: 7A65, subunit A), PTCH1 (PDB ID: 6RVD, subunit A) and PC2 (PDB ID: 6T9N, subunits A-D). Proteins are shown in white for P-gp/PTCH1 and grey for PC2 to differentiate subunit A from the other homotetrameric subunits of PC2. Cholesterols bound to Site A (blue) and Site B (red) are shown and the 6 site residues are indicated as cyan/pink spheres.

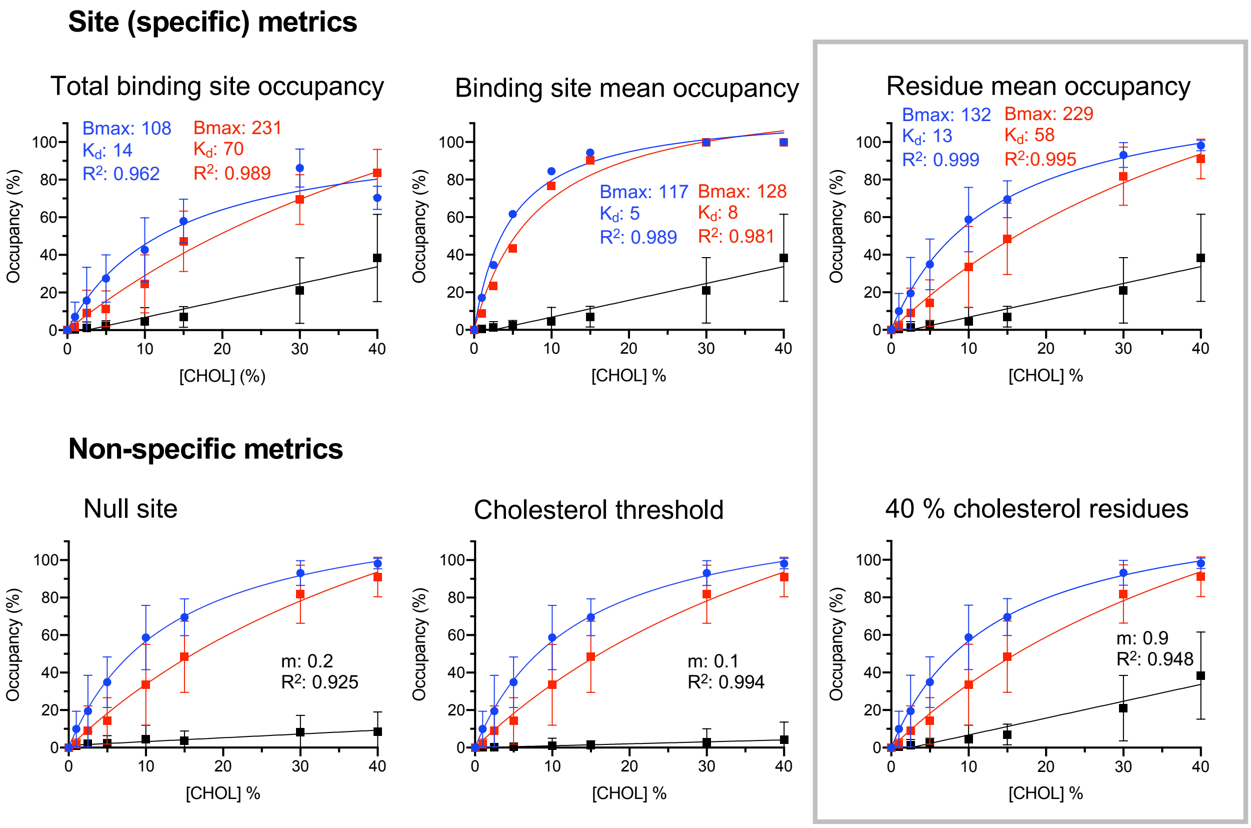


**Supplementary Figure 2: Methods for defining specific and non-specific interactions from the PyLipID outputs, exemplified using PC2.**

Top: tested metrics used to define site interactions. These included the binding site occupancy as reported by PyLipID (total binding site occupancy), the mean per residue occupancy of all residues in the site (binding site mean occupancy) or the mean occupancy of 6 sites residues with conserved interactions across all % cholesterol (residue mean occupancy).

Bottom: tested metrics for characterising non-specific interactions. These comprised the mean occupancy of all residues which had occupancies > 0 % but which were not assigned to a binding site during community analysis (null site), residues with occupancies less than the % cholesterol in the simulation (cholesterol threshold) or residues which had an reported occupancies 30-50% in the 40% cholesterol simulations (40% cholesterol residues).

The grey box indicates the selected metrics used to define site and non-specific interactions in Fig. 2B/3B/4B/5A.

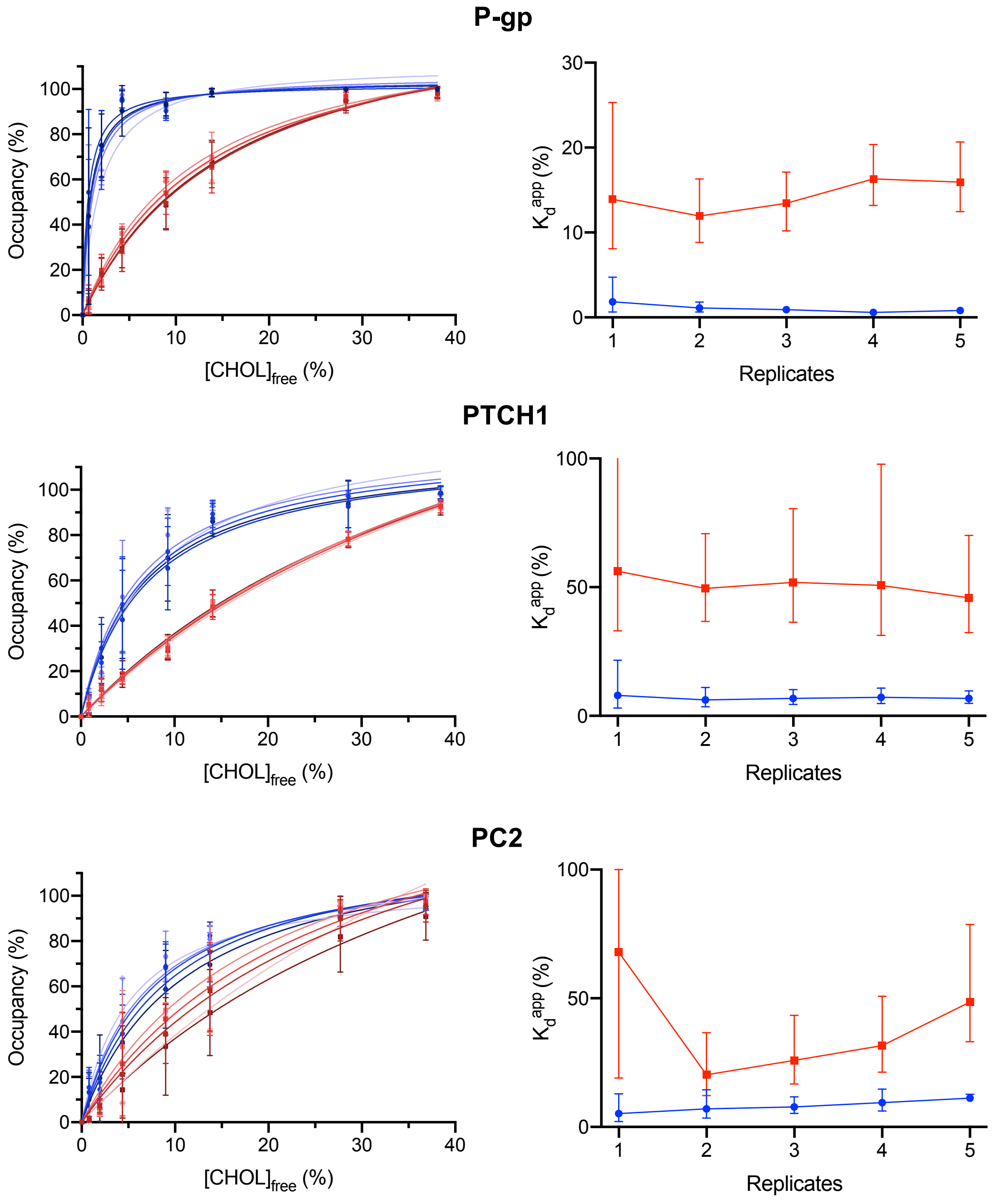


**Supplementary Figure 3: Binding saturation curve convergence.**
Left: Site A (blue) and B (red) binding saturation curves for P-gp, PTCH1 and PC2 when the number of replicate CG simulations was varied from 1 repeat (light blue/red) to 5 repeats (dark blue/red) at each % $[CHOL]_{free}$.
Right: Convergence of $K_d^{app}$ values as the number of replicate simulations was varied for Site A (blue) and Site B (red).

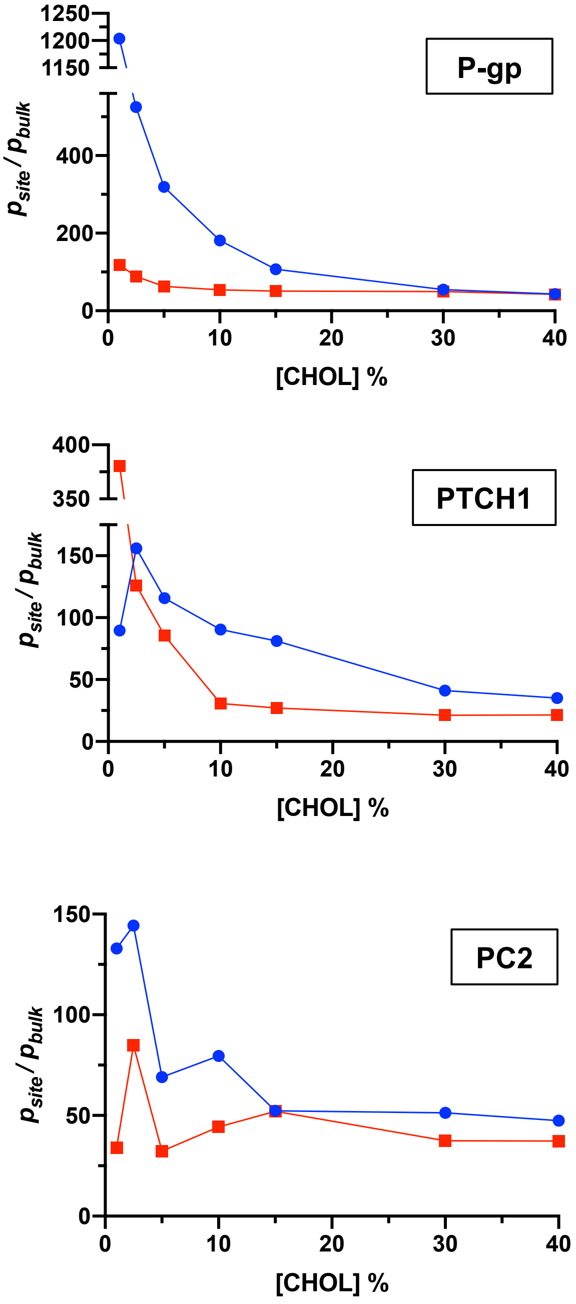


**Supplementary Figure 4: Probability of cholesterol localisation at a site from 3D density analysis.**

Probability of cholesterol localisation at Site A (blue) or Site B (red) ($p_{site}$) divided by the probability of cholesterol localisation within the bulk membrane region ($p_{bulk}$) from 3D density distributions of cholesterol across CG simulations. $p_{site}/p_{bulk}$ values were used to obtain free energies in Fig. 2D/3D/4D using Equation 3. Densities were calculated using MDAnalysis across 5 x 5 μs CG simulations at each % cholesterol. Further details of the density analysis are provided in the Supplementary Material.

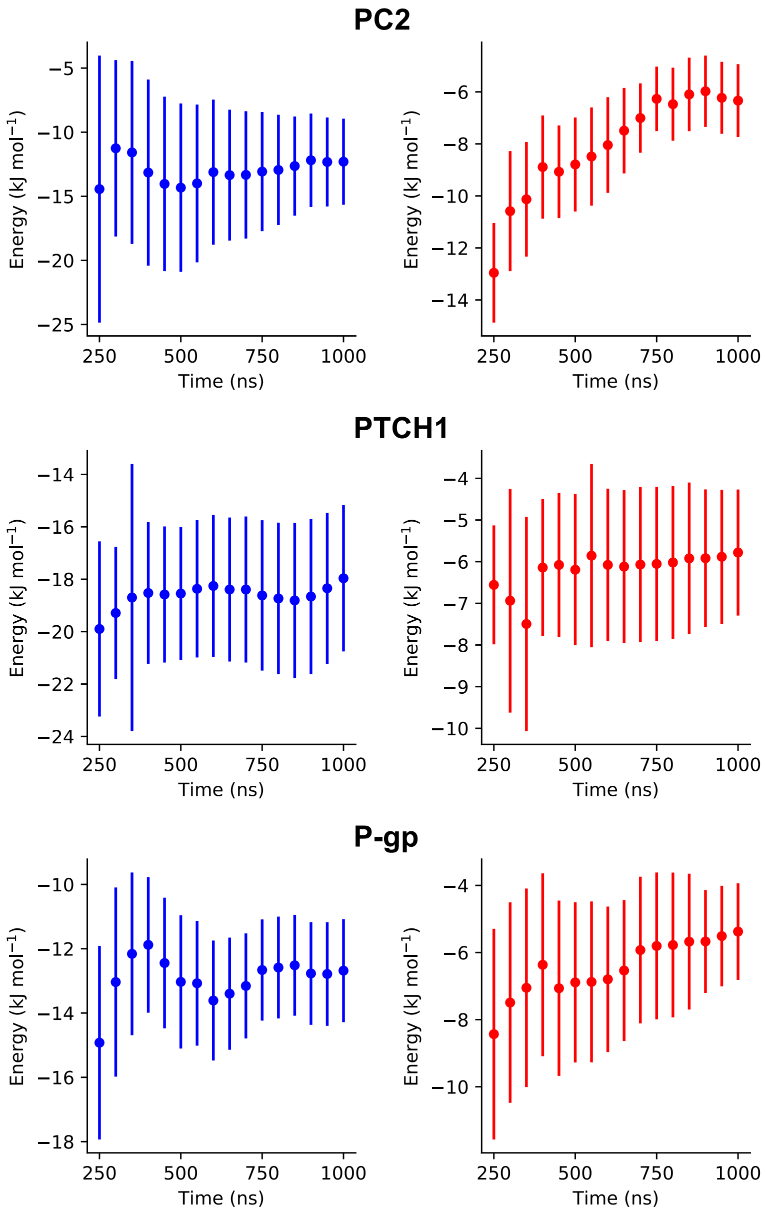


**Supplementary Figure 5: Convergence of site free energies derived from PMF calculations.**

Convergence of cholesterol binding Site A (blue) and B (red) free energy values derived from PMF calculations. The first 200 ns of each window was discarded and the window length was gradually increased in 50 ns intervals up to 1 μs.

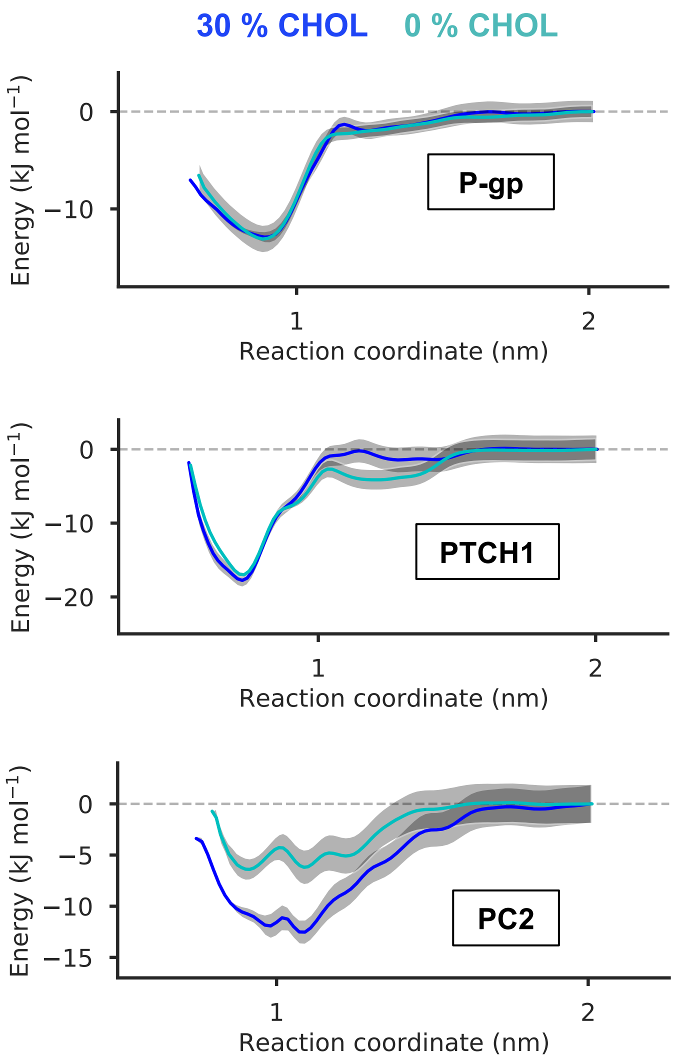


**Supplementary Figure 6: PMF profiles in the presence of differing amounts of membrane cholesterol.**

PMF profiles for cholesterol bound at Site A of P-gp, PTCH1 and PC2 when the total abundance of membrane cholesterol was 30 % (dark blue) or 0 % (i.e. only the site cholesterol was present, cyan).

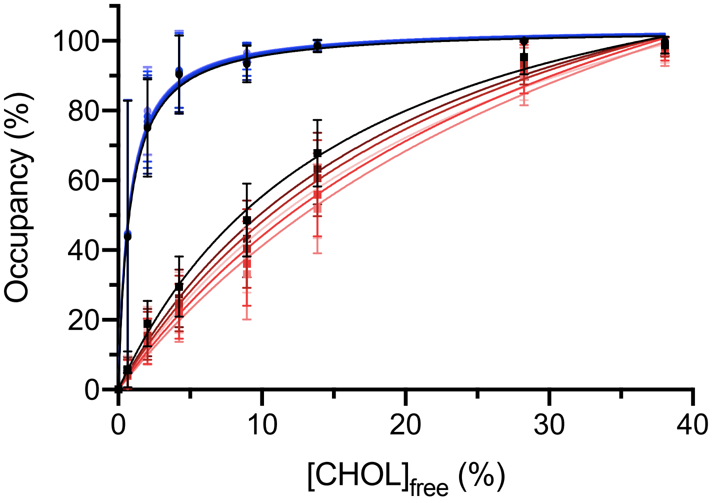


**Supplementary Figure 7: Binding saturation curve variability with number of site residues.**

Binding saturation curves for cholesterol binding to Site A (blue) or B (red) of P-gp. The number of residues used to define the binding site was varied from 1 (light blue/red) to 6 (dark blue/red) residues.
